# What we can and what we cannot see with extracellular multielectrodes

**DOI:** 10.1101/2020.12.21.423760

**Authors:** Chaitanya Chintaluri, Marta Bejtka, Władysław Średniawa, Michał Czerwiński, Jakub M. Dzik, Joanna Jędrzejewska-Szmek, Kacper Kondrakiewicz, Ewa Kublik, Daniel K. Wójcik

## Abstract

Extracellular recording is an accessible technique used in animals and humans to study the brain physiology and pathology. As the number of recording channels and their density grows it is natural to ask how much improvement the additional channels bring in and how we can optimally use the new capabilities for monitoring the brain. Here we show that for any given distribution of electrodes we can establish exactly what information about current sources in the brain can be recovered and what information is strictly unobservable.

We demonstrate this in the general setting of previously proposed kernel Current Source Density method and illustrate it with simplified examples as well as using evoked potentials from the barrel cortex obtained with a Neuropixels probe and with compatible model data.

**Author Summary:** Every set of measurements is a window into reality rendering its incomplete or distorted picture. It is often difficult to relate the obtained representation of the world to underlying ground truth. Here we show, for brain electrophysiology, for arbitrary experimental setup (distribution of electrodes), and arbitrary analytical setup (function space of current source densities), that one can identify distributions of current sources which can be recovered precisely, and those which are invisible in the system. This shows what is and what is not observable in the studied system for a given setup, allows to improve the analysis results by modifying analytical setup, and facilitates interpretation of the measured sets of LFP, ECoG and EEG recordings.

## Introduction

Multisite recording of extracellular potential is a popular technique in neuroscience. The potential obtained with a multitude of probes, from single wires, through silicone probes, CMOS arrays, to cortical ECoG and scalp EEG, reflects activity of underlying neural network and is directly related to the distribution of current sources along the active cells (current source density, CSD). The relation between the CSD and recorded potential, while occasionally contested (Bédard and Destexhe, 2011), overall is well established and trusted in experimental and analytical practice (Mitzdorf, 1985; Nunez and Srinivasan, 2006; Buzsáki, Anastassiou, and Koch, 2012; Einevoll et al., 2013; Gratiy et al., 2017). Due to the long range of electric potential and resulting high correlations between neighboring contacts (Łęski, Daniel K Wójcik, et al., 2007; Hunt et al., 2011; Lindén et al., 2011; Łęski, Lindén, et al., 2013) it is useful to estimate the current sources which can remove or decrease these spurious correlations.

Several methods have been introduced to estimate current sources since 1950s (Pitts, 1952; Nicholson and Freeman, 1975; Pettersen et ah, 2006; Łęski, Daniel K Wójcik, et al., 2007; Łęski, Pettersen, et al., 2011; Potworowski et al., 2012). Here, using the kernel CSD method (kCSD, Potworowski et al. (2012)), we show explicitly which sources are accessible experimentally for a given setup (distribution of electrodes and analytical assumptions) and which sources are strictly inaccessible (invisible) experimentally. We also show how using expert knowledge one may gain access to these otherwise invisible sources. We illustrate these new advancements with simplified examples as well as evoked potentials recorded with a Neuropixels probe (Jun et al., 2017) in the barrel cortex and with compatible model data. The Neuropixels probe is an example where a traditional approach to CSD estimation fails while it falls very naturally into the framework of kCSD.

### Current source density and extracellular potential

The extracellular potential that we measure is a consequence of ion motion in the tissue which is driven by ionic currents through the ion channels embedded in neuronal and glial membranes, as well as capacitive currents arising in response to potential gradients across the membrane. From the perspective of extracellular medium it seems as if the current was disappearing or appearing from inside a cell, which is why we talk about current sources and sinks. The distribution of these current sources is called the current source density (CSD) and its relation to the extracellular potential is given by the Poisson equation

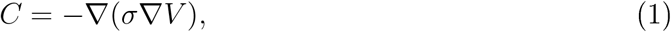

where *C* is the CSD, *V* is the extracellular potential, and *σ* — the conductivity tensor. Thus, if we knew the potential in the whole extracellular space, we could easily compute the CSD. On the other hand, knowing CSD in the whole space, we can compute the extracellular potential. Assuming isotropic and homogeneous tissue the Poisson equation reduces to

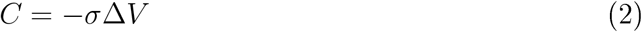

which can be easily solved:

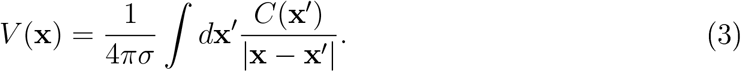

In more complex situations when σ depends on position and direction, and we have nontrivial boundary conditions, one must resort to numerical integration (Ness et al., 2015; Næss et al., 2017). Careful discussion of the meaning of the CSD and derivation of the relations between CSD and the potential can be found in Stevens (1966), Nicholson (1973), and Gratiy et al. (2017). Discussion of physiological sources of the extracellular potential can be found in the reviews by Buzsáki, Anastassiou, and Koch (2012) and Einevoll et al. (2013).

### Kernel Current Source Density estimation method

Kernel CSD estimation (Potworowski et al., 2012) is a two-step procedure. First, one does a kernel interpolation of the measured potential which gives *V*(*x*) in the whole space. This is obtained with the help of a symmetric kernel function, *K*(x, x′), so that

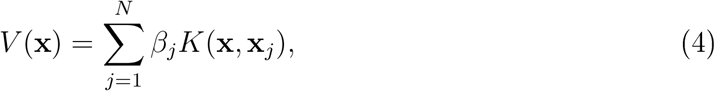

where x_*j*_ *j* = 1,…, *N*, are electrode positions. The regularized solution, which makes correction for noise, is obtained by minimizing prediction error

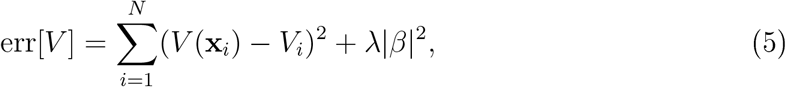

which gives

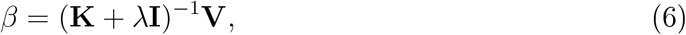

where **V** is the vector of measured potentials, λ is regularization parameter, and

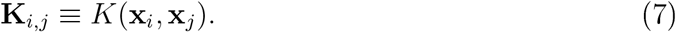

Once the potential is estimated the obtained solution must be moved to the CSD space. This is easiest to understand in 3D where one can simply plug the estimated potential into the Poisson equation [1], and compute CSD everywhere. In the general case this can be achieved with a second function, which we call cross-kernel, 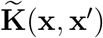. With these functions the resulting CSD estimation is given by

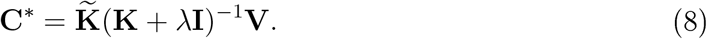

In principle one could consider arbitrary smoothing kernels **K** but in general it is then difficult to identify the relevant corresponding 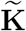, except in simple unphysiological cases such as infinite space with homogeneous conductivity. To simplify computation and improve understanding of the estimation space we introduce a large basis of CSD sources spanning the region of interest, 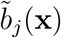, and corresponding basis in the potential space, *b_j_*(**x**), and construct our kernels from these basis functions (Potworowski et al., 2012). Thus the CSD and the potential are represented as

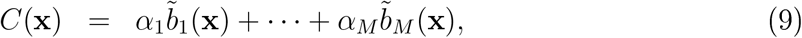

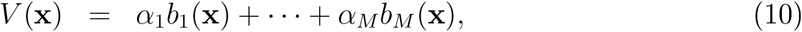

while the kernel functions are given by

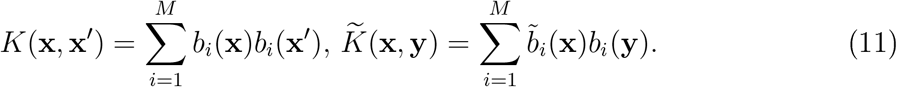

We recall the details of kCSD method in the Methods section.

## Results

Here we investigate how the experimental setup (distribution of electrodes), which is finalized during the experiment, and the analytical setup (distribution of basis functions), which can be varied post-experiment, affect reliability of the estimation. We show what can and what cannot be inferred about the neural activity in the brain for the selected combination of experimental and analytical approaches using a simple metaphor as well as model and experimental data from Neuropixels recording of somatosensory evoked potential in the barrel cortex.

### Spectral decomposition for kCSD and regularization

Let us reconsider the construction of kCSD (see Methods or Potworowski et al. (2012)). In kCSD we estimate CSD in space 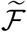 span by a large, *M*-dimensional basis, 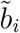. However, our experimental setup imposes constraints which force our model to an *N*-dimensional subspace where the estimation really takes place, with *N* ≪ *M*. To understand the structure of this smaller space we can decompose the operator **K**, eq. [7], acting on the measurements. We can take advantage of the symmetry and positivity of **K** matrix which guarantee existence of eigendecomposition

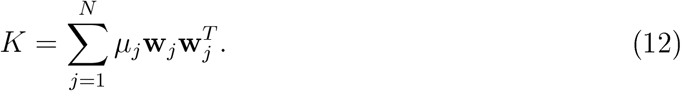

Then the kCSD reconstruction for a set of measurements **V**, eq. [8], is

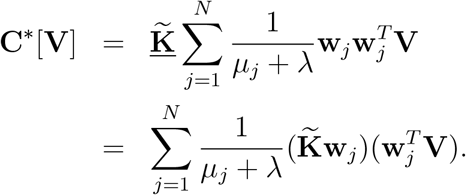

Since Wj are orthogonal we have

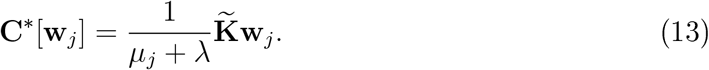

Thus **w**_*j*_ are the natural ‘eigenmeasurements’ corresponding to individual CSD profiles, 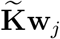, accessible to the given setup when specific basis 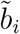 is assumed. Moreover, it is easy to see that the CSD profiles

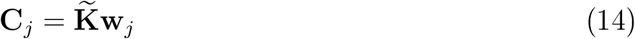

actually form the basis of estimation space, we call them ‘eigensources’, and we denote the space they span by 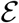. Since kCSD method is self-consistent, in the absence of noise, we see that the potential at the electrodes generated by **C**_*j*_ is

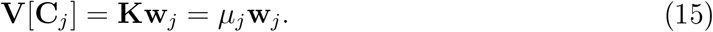

It leads to reconstructed CSD

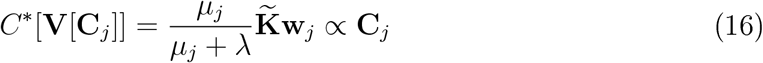

which is equal to **C**_*j*_ for λ = 0.

A natural question appears as to what happens to the missing *M* – *N* dimensions. The answer is that they are projected onto 0 (annihilated). It is possible to construct this space explicitly. Starting with the basis of *N* eigensources we can expand it within 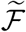 by Gram-Schmidt orthogonalization. This construction splits the space of all CSDs, 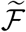, into two orthogonal subspaces, one of which spans all the sources which can be recovered with a given setup, 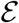, and its orthogonal complement, 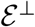, containing all the sources which are annihilated.

In Figure 1 we illustrate these concepts using a unit interval [0,1] as a metaphore for the whole brain. We consider a 1D Gaussian source centered at 0.25 (“left hemisphere”), which is essentially nonzero on interval [0, 0.5], In the left column we show the true source (ground truth, GT) and the kCSD reconstruction as well as the location of the electrodes where the potential was “measured”. The second column shows projections of the true source on the space of eigensources: this is the part of the true source accessible to the kCSD method. The third column shows the part of the true source which is annihilated, in other words, not accessible experimentally. We also show the difference between the ground truth and the kCSD reconstruction. The fourth column shows the eigensources for the given situation as a reference.

**Figure 1:**
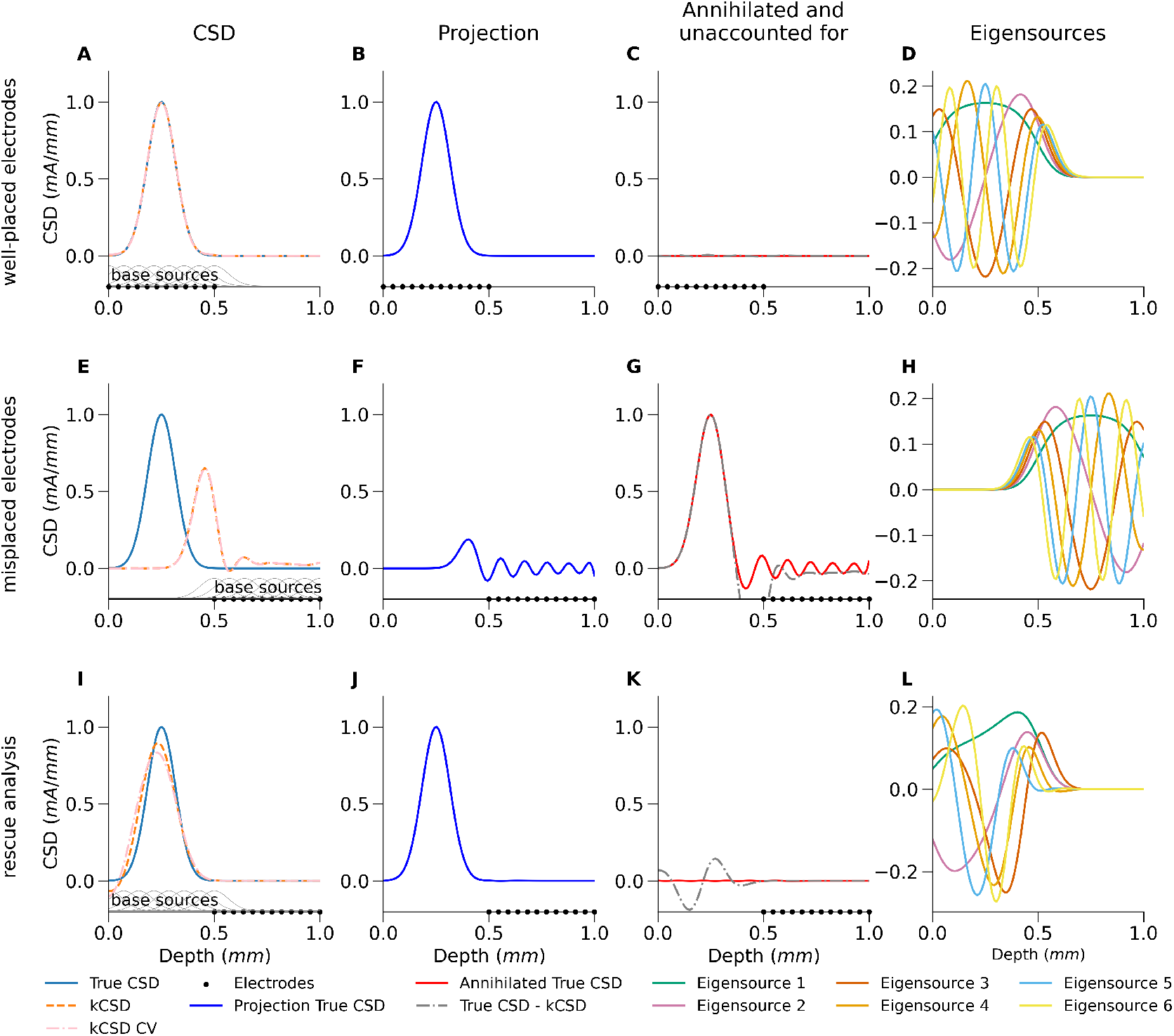
Proper placement of base sources allows analysis of data collected with misplaced electrodes. Demonstration of kCSD reconstruction for different relative placement of basis sources and electrodes for simple 1D Gaussian source in the absence of noise. Width of the basis source, *R*, was selected through cross-validation for the first case (A) and used throughout. Left column shows the true source (continuous line) and the kCSD reconstruction (broken line) as well as the location of the electrodes where the potential was “measured” (dots on the horizontal axes) and the positions of the basis sources (gray Gaussians on lower axis, first column). Second column: projections of the true source on the space of eigensources, the part of the true source accessible to the kCSD method. Third column: the part of the true source which is annihilated or not accessible experimentally and the difference between the True CSD and the reconstruction (broken line). Fourth column: the eigensources for the given setup. First row: the electrodes and the base sources are placed in the region containing the source to be reconstructed ([0, 0.5]). Second row: the electrodes were badly misplaced ([0.5,1]) and the base sources placed where the electrodes are (standard CSD analysis). Third row: the electrodes were badly misplaced ([0.5,1]), to compensate during analysis the base sources were placed where the source was expected, ([0,0.5]).

The first row shows the situation where the electrodes (dots on the horizontal axes) and the base sources (small gray Gaussians) were placed in the region containing the source to be reconstructed ([0, 0.5], “left hemisphere”). In this case the reconstruction is almost perfect as visualized by the almost zero difference between the reconstruction and the GT (panel B) and the annihilated part of GT being close to zero.

In the second row we show the situation where the electrodes were badly misplaced ([0.5,1], placed in the “right hemisphere”) and the base sources were placed where the electrodes are (standard CSD analysis). This example shows that we have one more aspect of the story we must discuss. In this case the projection of GT onto 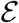 shown in panel F is much worse than the reconstructed kCSD, even though both are in 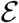. How is this possible? We can see that our space of current sources 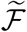 is an artificial construct since it is defined by the experimenter. Here we assumed that the sources lie in the “right hemisphere” while in fact they are in the “left hemisphere”. Hence the projection of GT onto 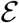 (panel F), unlike in the previous case, is so small mainly because the projection of GT on 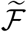 is small.

So to build a complete mental picture of the analytical situation in which we work in general observe that we have a universe of possible CSDs, let us call their space 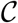. Then a given GT consists of two parts, one belonging to 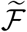 the other belonging to 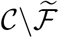. This second part cannot be reconstructed as it lies beyond the scope of our method. But it does generate measurable potential, as in the case shown in the second row of Fig. 1. The reason the kCSD reconstruction in panel E is better than the projection of GT shown in F is that the method tries to explain the remaining potentials with available mechanisms (using the eigensources) by a sort of aliasing known from Fourier analysis. In this case we obtain a satisfactory result, but in general this may result in artifacts which are difficult to distinguish from real sources.

The third row illustrates the situation where the electrodes were badly misplaced as before ([0.5,1]) but in attempt to compensate this during analysis, the base sources were placed where the source was expected ([0,0.5]). What this shows is that even if the electrodes are badly misplaced, if there is insight where to expect the source, the analysis can be rescued and a reasonable reconstruction can be obtained, even with noise (not shown). This is a consequence of the fact that kCSD estimation (8) depends on both the experimental setup (distribution of electrodes), fixed at the time of the experiment, and the analytical setup (distribution of basis functions), which can be varied. The separation of analytical from experimental setup allows to save data which would otherwise be wasted. Note that here the reconstruction is not perfect as indicated by nonzero difference between kCSD and GT in panel F. This is inadequacy of the method as the GT clearly belongs to 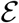 as indicated by zero annihilated part (panel G) but keep in mind we attempted reconstruction from data collected at very remote sensors.

This is a very important observation which we must emphasize: placing basis sources close to the activity, even away from the electrodes (panels E-H) may rescue the analysis recovering difficult data from misplaced electrodes or in cases where we do not cover all sources which we know contribute to the measurements. However, placing the basis sources where there is no activity may lead to methodic artifacts as outside sources may cast their shadows through the above aliasing mechanisms. Metaphorically, projections of GT on 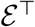 are known unknowns, while projections of GT on 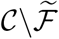 are unknown unknowns.

On a side, observe that since the experimental and analytical setups at the top and bottom rows are symmetric with respect to 0.5, the eigensources (panels D and H) are also symmetric. However, since their relation to the true CSD is different, the projections and annihilators are different.

### Example application: CSD reconstruction from Neuropixels probe

Here we illustrate the kCSD method in a realistic situation. We used data from simulated cortical recordings of whisker deflection in Traub’s model of a barrel column (Traub et al., 2005; Głąbska, Chintaluri, and Daniel K. Wójcik, 2016; Głąbska, Chintaluri, and Daniel K. Wójcik, 2017) with virtual sensors placed according to Neuropixels design as well as Neuropixels recordings from barrel cortex in a corresponding protocol. This example is interesting as it does not provide an obvious way to apply traditional CSD. The checker-board structure of a Neuropixels probe with only two contacts in each row alternating among four columns does not allow to compute directly traditional 2D CSD, that is apply numerical double derivative in rows. One could do it along diagonals ending up with two columns of CSD estimates alternating one point per row. Alternatively, one could compute 1D CSD along the four columns of the shank and further process, e.g. interpolating and averaging. The point is that any processing of this kind would be conceptually equivalent to a model-based CSD analysis, except the assumed rationale would be implicit and not clear. Kernel CSD and other explicit model-based analysis methods, such as iCSD, spell out the model explicitly, so one can investigate the role of one’s assumptions on the results. Further, kCSD, separating the reconstruction space from experimental setup gives a better control over analysis in such involved cases, as for Neuropixels probes.

We start with a model of thalamocortical loop (Traub et al., 2005; Głąbska, Chintaluri, and Daniel K. Wójcik, 2016; Głąbska, Chintaluri, and Daniel K. Wójcik, 2017). The full model we studied contains 3560 cells in multiple cortical and thalamic populations. In Fig. 2 we used one moment from a dataset `pulsestimulus10model.h5` we published previously (Głąbska, Chintaluri, and Daniel K. Wójcik, 2016; Głąbska, Chintaluri, and Daniel K. Wójcik, 2017), which contains recorded transmembrane currents from 10% of all cells. Only cortical contributions were used here. We simulated current injection to thalamic cells to induce cortical activity corresponding to a response to a whisker flick in the rat barrel cortex. Figure 2.A shows positions of all simulated cell segments we used from within a slice of 100 *μm* width passing through the center of the simulated barrel column. The color encodes the value of transmembrane currents. Figure B shows the True CSD computed in voxels of 50 *μm* side within a slice passing through the column center.

**Figure 2:**
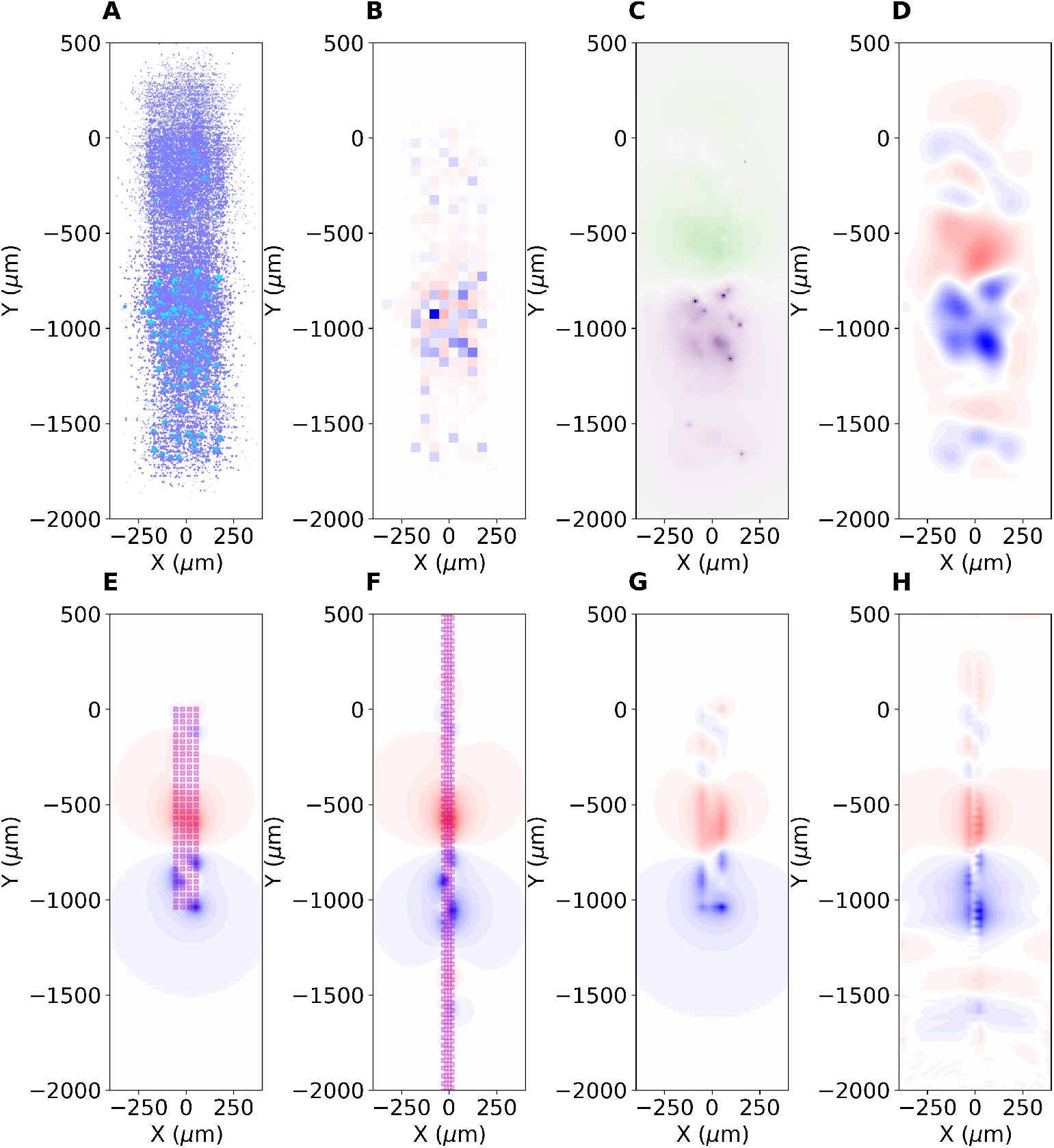
Validation of kCSD on data from simulated cortical recordings of whisker deflection in Traub’s model. (A) Colored dots indicate positions of simulated cell segments within a slice of 100 *μm* width passing through the center of the simulated barrel column. The color encodes the value of transmembrane currents. (B) True CSD computed in voxels of 50 *μm* side, slice passing through the column center. (C) Extracellular potential computed from the whole column in the place passing through the center of the column. (D) True CSD smoothed with a Gaussian kernel. (E, F) CSD estimated with kCSD with cross-validation from simulated measurements with a single electrode block of a Neuroseeker (E) and a single bank of Neuropixels (F) probes. (G) and (H): Projections of D on the space of eigensources defined by setups shown in (E) and (F). Magenta squares indicate the positions of electrode contacts.

These data allow calculation of extracellular potential (Głąbska, Chintaluri, and Daniel K. Wójcik, 2017). Figure 2.C shows the extracellular potential computed on the plane passing through the center of the column. Contributions from all the traced cells in the column were taken into account. Figure D shows the true CSD (Fig. B) smoothed with a Gaussian kernel. Figures E and F show CSD estimated with kCSD with cross-validation from simulations of potential measurements with modern probes. E) shows a reconstruction from a 4×32 probe, which follows the design of a single electrode block of the Neuroseeker (Raducanu et al., 2017), where we ignored the reference electrode. Figure F shows a reconstruction from simulated recordings with one bank of Neuropixels probe (Jun et al., 2017). Magenta squares indicate the positions of electrode contacts used in the computation. We assumed that both electrodes pass through the center of the column.

Observe that both probes seem to capture the dominating polarity changes across layers. Clearly, the broader span of the ‘Neuroseeker’ probe improves resolution within the layer, while the longer extent of the ‘Neuropixels’ probe better resolves the changes in higher layers. In both cases we can see that the features in space observed in CSD in Figure D are dragged towards the electrode by the reconstruction which should be taken into account in interpreting results from actual experiments. Note that in this analysis we placed the basis sources in the whole region represented on the picture. The explanatory power of these probes is shown in plots G and H. Here we show projections of the true source on the respective eigensources, which, as explained before, are defined by both the electrode setup and source placement. One can clearly see the specifics of the setup significantly affect the shape of the CSD part which can be reconstructed. In practice many elements would affect the reconstruction: true Neuroseeker probe has 12 times more channels available. Both probes would disrupt tissue and these effects are difficult to compare and go significantly beyond the scope of this illustratory section.

In figure 3. A and B we show the LFP and CSD profiles estimated along the column axis as functions of time for the same simulated dataset. Both figures 2 and 3.A and B clearly show that the model example used here is physiologically rather simple: the difference between the LFP and CSD profiles is minor. This is to be expected as we model only a single column, with two main cell populations contributing to the visible activity (Głąbska, Potworowski, et al., 2014). We can expect much more complex situations in actual experiments where multiple columns will contribute to the recorded activity, Figure 3.C and D. The broken vertical line in Figure 3.C indicates the time when Figure 5 was drawn. Note an additional dipole present superficially from −1mm, whose location is consistent with the putative microcolumn visible in Figure 5. Note that this dipole is difficult to discern in the LFP profile. Horizontal lines on the plot of CSD indicate the positions of layers based on histology (not shown). The visible stripy pattern seems to be a consequence of the probe design as discussed below.

**Figure 3:**
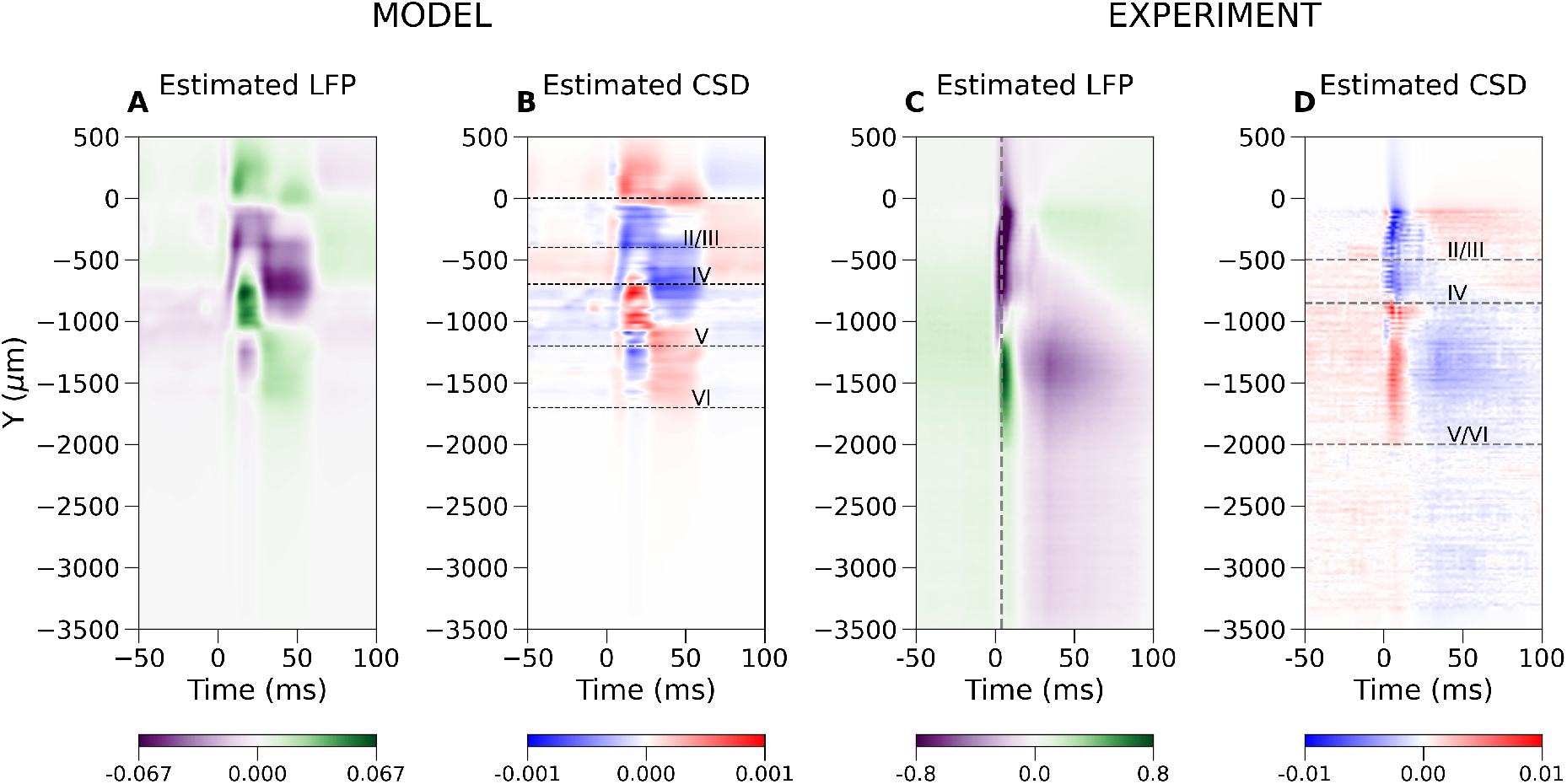
A. LFP and B. CSD within the model column as a function of time. Horizontal lines indicate position of the layers according to the model definition, which defines distribution of cell bodies. Corresponding C. LFP and D. CSD in actual experiment using Neuropixels probe in rat barrel cortex. Dashed vertical line in C marks the time snapshot where Fig. 5 is plotted.

**Figure 4:**
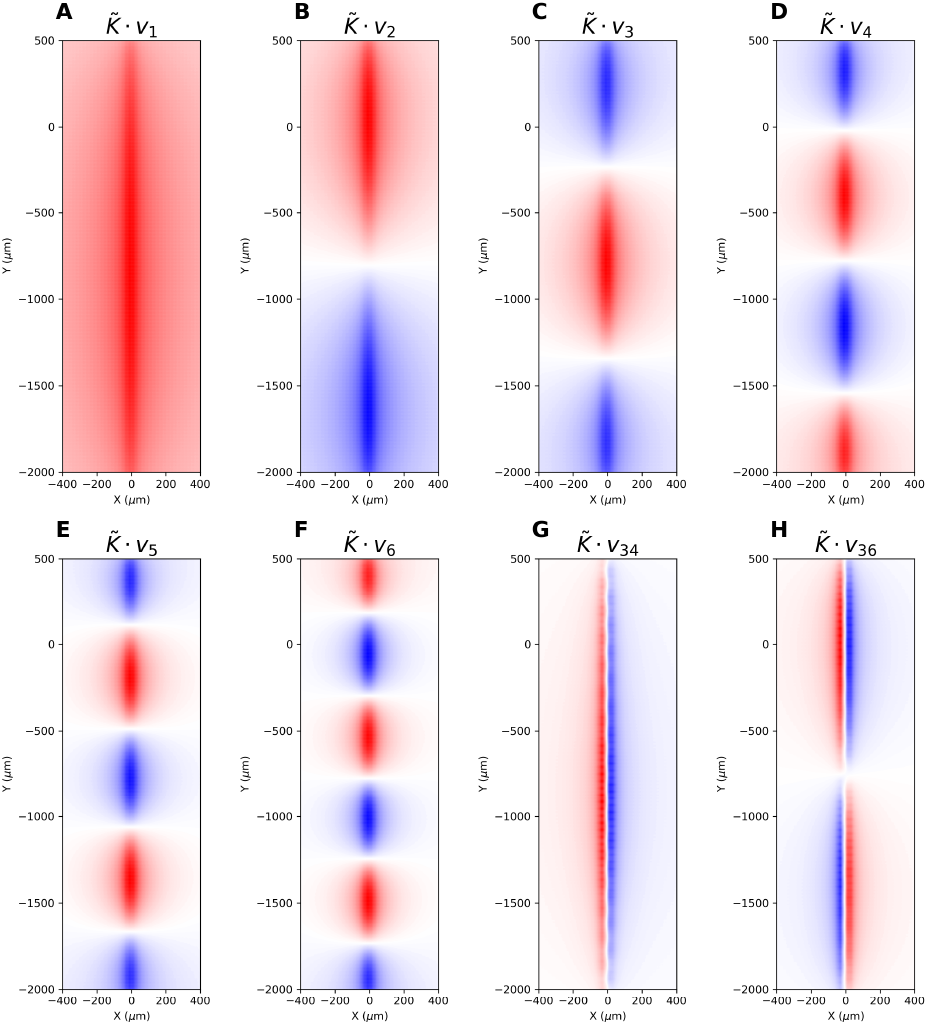
The first six eigensources and the first two eigensources spanning the horizontal dimension for a single bank of a Neuropixels probe.

**Figure 5:**
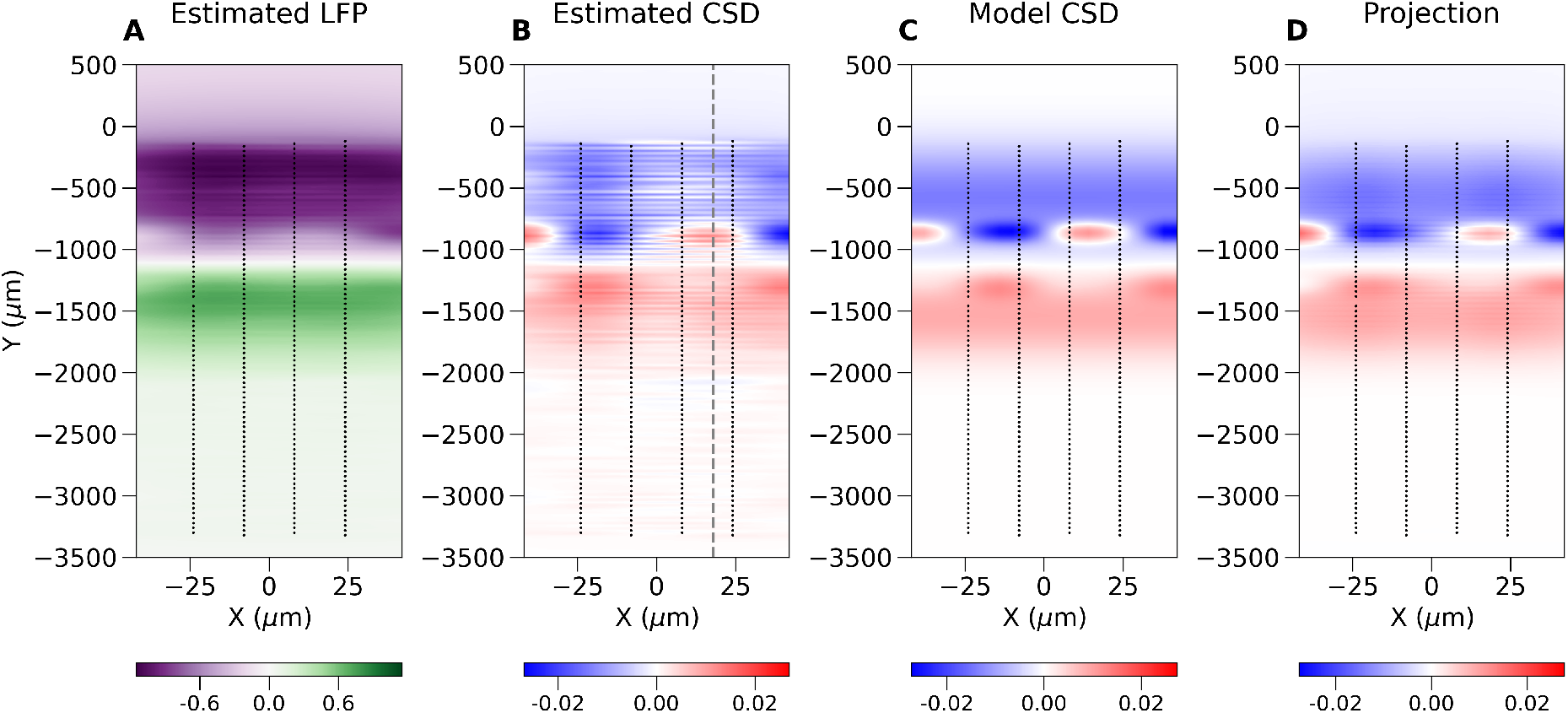
A. LFP and B. CSD profiles within the recording plane during the peak of response. The paired sinks and sources originating in layer Va (around −900 μm) could be the microcolumns within the barrel cortex macrocolumn. C. Speculated combination of five dipoles which could explain main part of the observed data seen in B. D. shows projection of C onto the space spanned by the eigensources. Close inspection shows the same stripy pattern visible in panel B indicating this is a feature of this probe design, not a bug of the method.

We can apply the same analytical machinery introduced before to the Neuropixels probes or to any other electrode setup, to obtain insight into the analysis of both simulated and experimental recordings. For example, the 33 leading eigensources for Neuropixels probes, due to its elongated design, are, loosely speaking, the 1D Fourier modes (Panels A to F, Fig. 4). There are eigensources spanning the orthogonal direction, which allows some spatial details to be recovered, however, the first two modes are number 34 and 36 (Fig. 4, G and H), which indicates these are rather fragile under noise.

Figure 5 shows an example reconstruction of the LFP and CSD in the plane of the electrodes from a snapshot in time from an experimental recording in the rat barrel cortex. The selected moment in time is marked with the dashed vertical line in Fig. 3.C. Black dots indicate positions of the electrodes. To make pictures more readable the distances on both axes differ strongly which introduced significant distortion visible in the figure. The vertical broken line in Fig. 5.B indicates location in space where data were taken for figure 3.C and D. We speculate that the paired sinks and sources (the blue and red spots) originating in layer Va could be the microcolumns within the barrel cortex (Buxhoeveden and Casanova, 2002). As indicated by the above analysis of model data, keep in mind that the features observed in the signal can actually represent sources placed more distally from the probe center, although the regularity observed in the position of those sources makes their estimated placement plausible. Note that these structures are difficult to discern from interpolated LFP, and hard to guess from direct LFP recording, which is available only at the points of contact. To support plausibility of this interpretation we postulated a model CSD profile consisting of four small and one extended dipoles built of pairs of gaussian sources and sinks (C). Their amplitudes were scaled to fit the estimated CSD profile (B). Next, we calculated eigensources for the recording setup and we projected the model CSD onto the space span by the eigensources (D). Although the reconstruction of the model CSD (not shown) for the studied probe was faithful, the same horizontal stripy artifacts visible in experimental analysis (panel B) could be observed, just as in projection (D). This shows that they arise due the Neuropixels probe design rather than as a bug of the method.

## Discussion

In this work we presented conceptual and computational tools to identify what can and what cannot be inferred from multielectrode recordings regarding the distribution of current sources which contributed to the measured potential. Using the kernel Current Source Density method (Potworowski et al., 2012) which separates analytical framework from experimental settings we showed how to identify which part of sources can be recovered and which is invisible. We introduced the concept of eigensources, eq. [14], which span the space of all possible solutions for a given experimental and analytical setup. They have the property that if the true CSD is one of the eigensources it can be recovered perfectly from noise-free measurements. We showed that the analytical space of all current sources we consider, 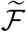, divides into two parts. One part is span by the set of eigensources, 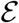, the other, 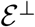, is annihilated, that is, any part of true CSD which belongs there does not contribute to the measured potential. Further, we showed that the analytical space we use is of necessity a subspace of all possible CSDs, 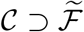, some of which do not belong to our analytical space 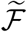 but may contribute nonzero potential (‘unknown uknowns’). An example is when the electrodes are in the thalamus, analytical space for reconstruction is in the forebrain, but strong source are in the cortex. In such a case we observe aliasing, where fake sources appear within the analytical space to compensate for the external sources. Finally, we showed how using expert insight one can improve source reconstruction by forcing it to the relevant region (Fig. 1).

We illustrated these concepts with simulated (Fig. 2 and 3 A and B) and experimental examples of cortical responses to rat vibrissae stimulation (Fig. 3 C and D and Fig. 5). We showed how one can combine simulation of current sources and extracellular potential with kCSD spectral analysis to understand which features of true sources can be recovered and to indicate possible artifacts in CSD analysis on specific probes. Importantly, we showed how this type of analysis can be applied data from modern probes, such as Neuroseeker or Neuropixels, for which traditional approaches to CSD estimation cannot be directly applied. Using experimental recordings with Neuropixels probe we showed that kCSD can significantly improve contrast and enhance certain features of ongoing activity in challenging situations, such as significantly different number of electrodes in vertical versus horizontal directions. Interestingly, despite having only 2 electrodes in each row of Neuropixels, in layer Va we were able to identify robust features (Fig. 3 D and Fig. 5 B) which we speculate are the microcolumns within the barrel cortex macrocolumn (Buxhoeveden and Casanova, 2002).

Finally, we note that although the analysis here has been presented for kernel CSD method, the kernel approach of Potworowski et al. (2012) can be adapted to other measurement modalities and experimental contexts, where the measurements are a function of underlying activity. Natural examples include other electrophysiological measurements, such as EEG, ECoG, SEEG, where one naturally combines geometry and conductivity of the brain and skull to improve precision. We currently develop the necessary tools to handle these measurements for animal and human brains. We are convinced that one can go further to other modalities, such as all types of calcium or voltage sensitive dyes, magnetic resonance, and others. This will be the subject of further study.

## Methods

### Experimental methods

An electrophysiological experiment was performed on an adult male Wistar rat (480 g). Procedures followed the 2010/63/EU directive and were accepted by the 1st Warsaw Local Ethics Committee. The animal was anaesthetized with urethane (1.5 mg/kg, i.p., with 10% of the original dose added when necessary) and placed in a stereotaxic apparatus (Narishige Group, Japan). A local anesthetic (Emla, 2.5% cream, AstraZeneca) was applied into the rats’ ears and the skin over the skull was injected with a mixture of Lignocaine (0,5%) and Bupivacaine (0.25%, Polfa Warszawa S.A.) prior to the surgery. Fluid requirements were fulfilled by s.c. injections of 0.9% NaCl. The body temperature was kept at 37°C by a thermostatic blanket.

The skull was opened to give access to the right barrel field (AP 1.5–2.5 posterior to Bregma, L 5–6 mm from middle line). A Neuropixels probe (imec) mounted on a micro-manipulator was inserted into the brain approximately perpendicular to the surface (30 degree from vertical axis) aiming through a barrel cortex to the somatosensory thalamus. Presented data was obtained from the probe inserted to the depth of 3500 *μ*m (Fig. 6).

**Figure 6:**
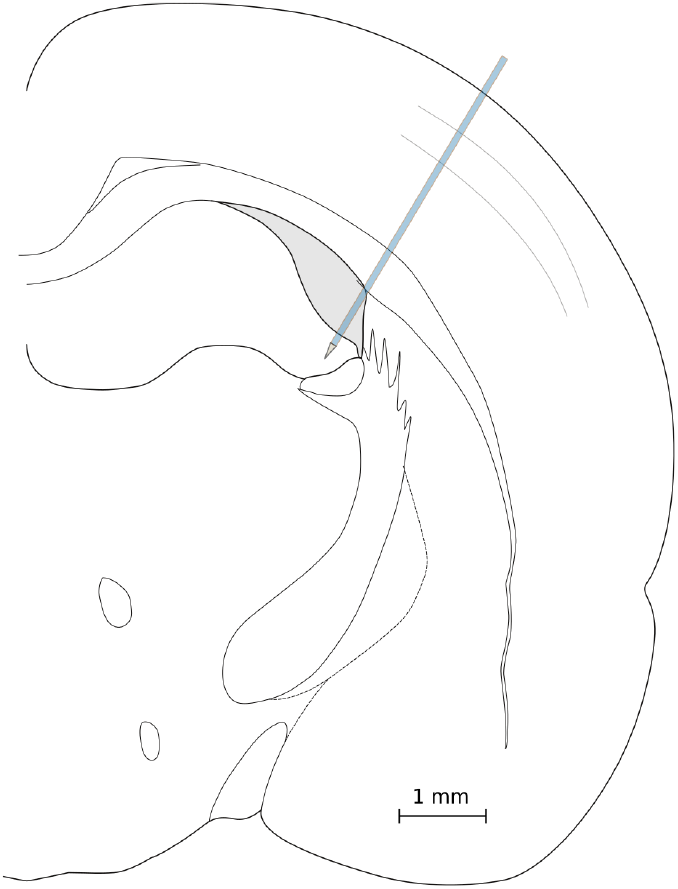
Location of Neuropixels bank 0 on schematic outline of a rat brain at the level of the barrel cortex (after Paxinos and Watson (2007)).

After completion of the experiment, the rat received an overdose of urethane and was perfused transcardially with phosphate buffered saline (PBS) followed by 10% formalin in PBS. The brain was removed and cryoprotected in 30% sucrose solution. Coronal sections (50 *μ*m) were cut on a freezing microtome for microscopic verification of electrode position.

To activate a barrel column in the right somatosensory cortex, the left C1 whisker was glued to a piezoelectric stimulator at around 10 mm from the snout. Spike2 software and CED 1401power interface (Cambridge Electronic Design, UK) controlled square pulses (1 ms, 25 V) that produced a 0.15 mm horizontal (in the rostro-caudal axis) deflection of the whiskers. The stimuli were delivered with 3–4 s pseudorandom intervals. Stimulation time stamps were recorded with digital input channels of a National Instruments card synchronized with the imec system.

Wideband signal was acquired with Neuropixels probe (v. 1.0, imec, Jun et al 2017) and SpikeGLXsoftware (v. 20190413-phase3B2, https://billkarsh.github.io/SpikeGLX/) using x500 gain and 30 kHz sampling rate. The data was acquired from all 384 electrodes in bank 0 (spanning 3840 *μ*m), arranged on four-column checkerboard configuration as in Figure 2.F. 64 electrodes extending above the cortex were discarded from the analysis shown here, so only 320 electrodes were used. Before analysis the data was filtered offline (1–300 Hz bandpass) and downsampled to 2.5 kHz.

### Review of Kernel Current Source Density estimation

#### Basis functions

For ease of reference here we repeat the key steps in the construction of kCSD estimation framework (Potworowski et al., 2012) to introduce the notation and establish the basic notions. We assume potential *V*_1_,…, *V_N_* was measured with ideal electrodes at points **x**_1_,…, **x**_*N*_.

We first construct a pair of related function spaces in which we perform the estimation, space of current sources 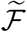 and space of potentials 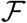, which are built of *M* ≫ *N* basis functions *b_i_* and 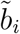

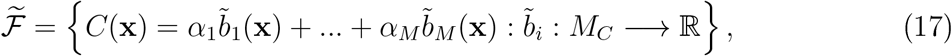

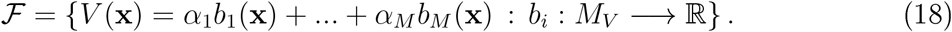

We select the basis source functions 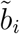 so that they are convenient to work with, such as step functions or gaussians, with support over regions which are most natural for the problem at hand. For example, when reconstructing the distribution of current sources along a single cell from a set of recordings with a planar microelectrode array, *M_C_* is the neuronal morphology, which we take to be locally 1D set embedded in real 3D space, while *M_V_* would be the 2D plane defined by the MEA.

The potential basis functions, *b_i_* are defined as the potential generated by 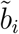, so that 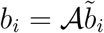, where 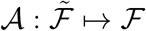. Specific form of 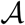 operator depends on the problem at hand, the dimensionality of space in which estimation is desired, as well as on physical models of the medium, such as tissue conductivity, slice or brain geometry, etc. (Pettersen et al., 2006; Łęski, Daniel K Wójcik, et al., 2007; Łęski, Pettersen, et al., 2011; Ness et al., 2015; Cserpan et al., 2017). In the simplest case of infinite, homogeneous and isotropic tissue in 3D we have

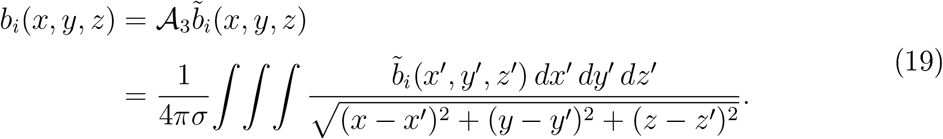

In general, we can consider arbitrary conductivity and geometry of the tissue which may force us to use approximate numerical methods, such as finite element schemes. For example, Ness et al. (2015) show an application of kCSD for a slice of finite thickness and specific geometry, as well as a method of images approximation for kCSD for typical slices on multielectrode arrays (recordings far from the boundary, slice much thinner than its planar extent).

In the past we considered CSD reconstruction for recordings from 1D, 2D and 3D setups under assumption of infinite tissue of constant conductivity (Potworowski et al., 2012), we used method of images to improve reconstruction for slices of finite thickness on MEA under medium of different conductivity (ACSF, Ness et al. (2015)) and we considered reconstruction of sources along single cells when we have reasons to trust the recorded signal to come from a specific cell of known morphology (Cserpan et al., 2017), where details can be found.

#### kCSD framework

We can think of all these potential basis functions *b_i_*(**x**) as features representing **x** in a *M*-dimensional space through related embeddings

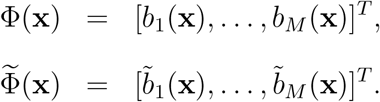

Let us introduce a kernel function in 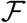 through

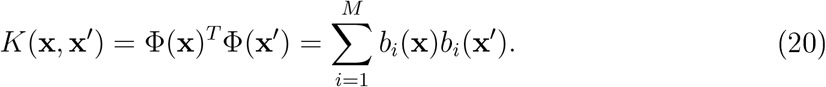

This kernel turns 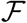 into a reproducing kernel Hilbert space (RKHS, Aronszajn (1950)) whose properties we discussed in Potworowski et al. (2012). In particular, we can show that all potential profiles admissible by our construction can be written as linear combinations of multiple kernels fixed with one leg at different points:

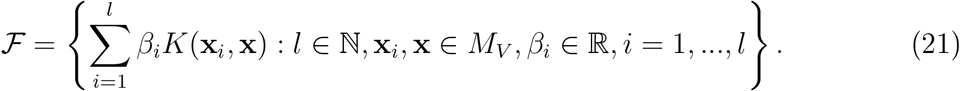

So we have now two representations of every function in 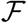, as a sum of kernels or a sum of basis elements

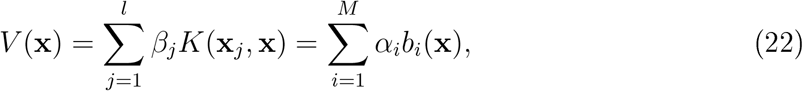

where

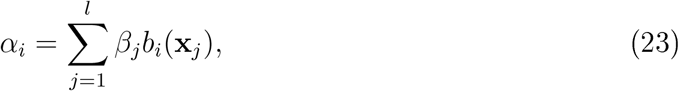

where **x**_*j*_ are some positions in space. One can see that the RKHS norm of such a potential function (Potworowski et al., 2012) in the two representations is

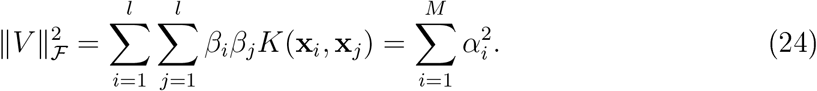

#### Source estimation with kCSD

Estimation of current sources with kCSD consists of two steps. The first is kernel interpolation of the potential, the second is changing the space from potential to sources. Conceptually, in the simplest case, this is equivalent to applying Laplacian or double derivative to the potential field obtained in the whole space. However, using our approach with double kernels, which take into account underlying physics and geometry of the studied system, it is possible to apply these ideas to more complex situations, e.g. slices of specific shape and conductivity profile (Ness et al., 2015) or fields generated by individual cells (Cserpan et al., 2017).

To estimate the potential in the whole space we minimize an error function

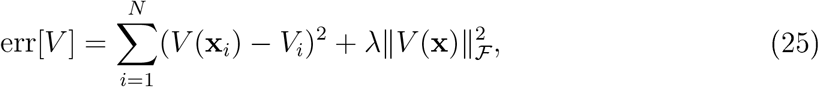

where the first term indicates proximity of our model to actual measurement, while the second constrains complexity of the model (Note a typo in Eq. (4.2) in the original paper, which incorrectly states the error term as 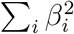 while it should be 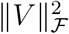, as given here by Eq. 24.). Using the representer theorem (Kimeldorf and Wahba, 1971) we can show that the solution is of the form

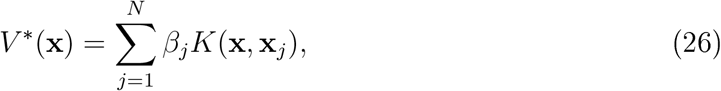

where **x**_*j*_ are the N electrode positions. Minimum of (25) is obtained for

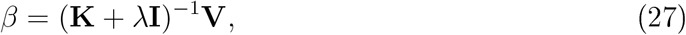

where

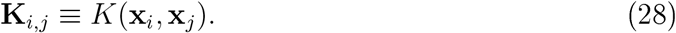

Now that we have the potential given by combination of kernels, Eq. (26), we can expand it in the original basis *b_i_*(**x**), Eq. (22). From that we obtain a consistent estimate of the CSD by lifting the model from the potential to the CSD representation:

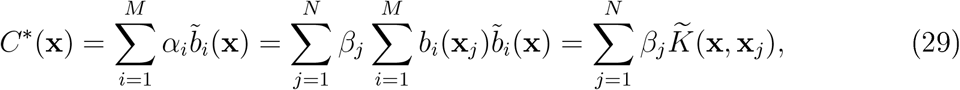

where we introduce the cross-kernel function (Note that this definition replaces the two variables with respect to the original definition from (Potworowski et al., 2012) to avoid transposition in the matrix formulation below.)

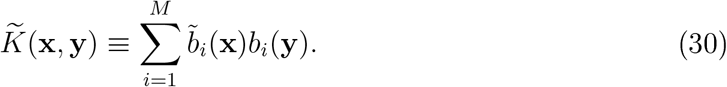

With this definition we can write

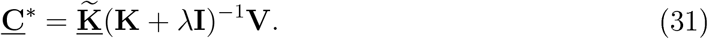

## Acknowledgments

The study received funding from the Polish National Science Centre’s grants (2013/08/W/NZ4/00691) and (2015/17/B/ST7/04123). K.K. is funded from European Research Council Starting Grant (H 415148; principal investigator: Ewelina Knapska). The authors declare no conflict of interest.

